# Molecular neuroanatomy of anorexia nervosa

**DOI:** 10.1101/440313

**Authors:** Derek Howard, Priscilla Negraes, Aristotle N. Voineskos, Allan S. Kaplan, Alysson Muotri, Vikas Duvvuri, Leon French

## Abstract

Anorexia nervosa is a complex eating disorder with genetic, metabolic, and psychosocial underpinnings. Using unbiased genome-wide methods, recent studies have associated a variety of genes with the disorder. We characterized these genes by projecting them into aggregated gene expression data from reference transcriptomic atlases of the prenatal and adult human brain. We found that genes from an induced stem cell study of anorexia nervosa are expressed at higher levels in the lateral parabrachial and the ventral tegmental areas. The adult expression enrichment of the lateral parabrachial is confirmed with genes from two independent genetic studies. In the fetal brain, enrichment of the ventral tegmental area is also observed for the six genes near the only common variant associated with the disorder (rs4622308). We also observed signals in the adult and fetal pontine raphe, but they were not observed when using the genes from the genetic studies. In addition to signals related to calcitonin gene-related peptide neurons and the tachykinin, we found more than the expected number of microglia marker genes within the gene sets. Using mouse transcriptomic data, we identified several anorexia nervosa associated genes that are differentially expressed during food deprivation. While these genes that respond to fasting are not enriched in the gene sets, we highlight *RPS26* which is proximal to rs4622308. We did not observe expression enrichment in the cingulate cortex or hypothalamus suggesting other targets for deep brain stimulation should be considered for severe cases. This work improves our understanding of the neurobiological causes of anorexia nervosa by suggesting disturbances in subcortical appetitive circuits.

## Introduction

Anorexia nervosa is a complex eating disorder that primarily occurs in women with onset often during adolescence. Currently, there are no effective treatments that can reverse the self-induced starvation, disturbed body image, and fear of weight gain that define the disorder (Watson and Bulik 2013). Patients often relapse and it is often considered to be a chronic condition (Zipfel et al. 2000). Due to self-starvation and suicide, it has the highest mortality rate of any psychiatric illness (Sullivan 1995).

Our understanding of the neuroanatomical circuits involved in anorexia nervosa is limited. MRI studies of people affected with the illness are limited to examining large structures, finding broad volume reductions and white matter alterations (Phillipou, Rossell, and Castle 2014; Miles et al. 2018). Diffusion tensor imaging studies have found alterations in several white matter tracts (Martin Monzon et al. 2016). Due to coarse resolution, MRI studies cannot detect structural or functional changes at the microcircuit level. In contrast, animal experiments have deciphered subcortical neurocircuits that control appetite and feeding behaviours. We note that these animal experiments are focused on true anorexia (loss of appetite) and do not mimic the complex symptoms of anorexia nervosa. Nonetheless, the neural circuits found in mice may provide therapeutic targets for patients with the disorder.

Importantly, anorexia nervosa has a strong genetic basis. Twin-based heritability is estimated to range from 48% to 74% (Yilmaz, Hardaway, and Bulik 2015) and genetic studies are starting to identify candidate genes. Specifically, three past studies have used genome-wide scans to associate genes and genetic variants with the disorder. First, a transcriptomic study that compared induced neural stem cells of anorexia nervosa patients and controls identified hundreds of differentially expressed genes (Negraes et al. 2017). Second, a genome-wide association study has reported a single significant locus for anorexia nervosa (rs4622308) that overlaps with six genes (Duncan et al. 2017). Finally, a study that performed whole exome sequencing identified damaging rare variants that are associated with disordered eating (Lutter et al. 2017). These genes do not provide a direct link to a specific brain region or cell type because they were derived from fibroblasts or DNA. In addition, it’s unknown if these genes respond to fasting. Using several transcriptomic resources, we characterized the anorexia nervosa associated genes across these dimensions (Figure 1).

**Figure 1.**
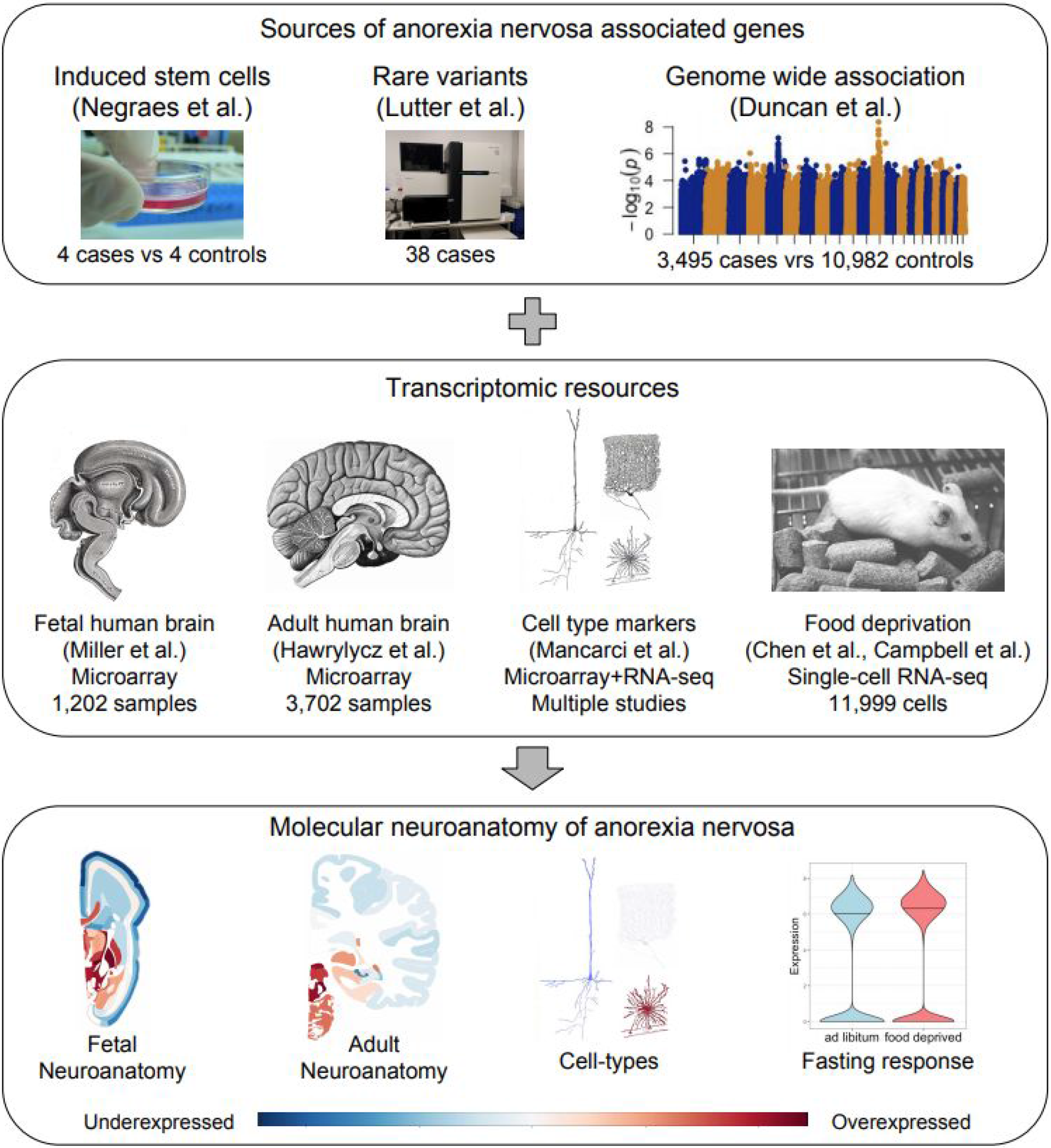
Overview of the study. Genes associated with anorexia (top) are characterized in several genome-wide expression datasets (middle) to provide locations and cell types (bottom). Images are from the cited publications, and Wikimedia Commons (Gray’s Anatomy by Henry Vandyke Carter; users kaibara87, Rama, and Konrad Förstner).

## Methods

### Adult human brain gene expression data

The Allen Human Brain Atlas provides a comprehensive transcriptional landscape of six normal human brains (Hawrylycz et al. 2012). Complete microarray gene expression datasets were downloaded from the Allen Human Brain Atlas data portal (http://human.brain-map.org/static/download/). These datasets were obtained from six individuals (five males, one female), with age ranging from 24 to 57 years. Custom 64K Agilent microarrays were used to assay genome-wide expression in 3,702 spatially-resolved samples (232 named brain regions). The Allen Institute normalized the data with a multistep process that adjusted for array-specific biases, batch, and dissection method. Full details of the procedures used by the Allen Institute researchers are available in the Allen Human Brain Atlas technical white paper (http://help.brain-map.org/display/humanbrain/Documentation).

### Prenatal human gene expression data

Similar to the Allen Human Atlas, this transcriptomic atlas of the normal mid-gestational human brain provides a brain- and genome-wide reference (Miller et al. 2014). Complete microarray gene expression datasets for were downloaded from the Brainspan website (http://www.brainspan.org/static/download.html). Datasets were obtained from 4 intact mid-gestational human brains that passed several exclusion criteria (15-21 postnatal weeks, 3 females). The same custom 64K Agilent microarrays that were used for the adult atlas were used to assay expression in the 1,203 spatially-resolved samples (516 named brain regions). Details of the procedures used by the Allen Institute researchers are available in the Brainspan Atlas of the Developing Human Brain technical white paper (http://help.brain-map.org/display/devhumanbrain/Documentation).

### Microarray gene expression data processing

Microarray probes were re-annotated using the Re-Annotator mRNA reference annotations to increase the total number of annotated probes (Arloth et al. 2015). For each donor, samples mapping to the same named brain region were mean averaged to create a single expression profile for each region. Analogous named brain regions of both hemispheres were not distinguished because no significant differences in molecular architecture have been detected between the left and right hemispheres (Miller et al. 2014). Expression levels of the 58,692 probes were then summarized by mean averaging for each of 20,869 gene transcripts. Gene expression values were next converted to ranks within a named brain region and z-score normalized across brain regions. Z-scores for each donor were then averaged across donors to obtain a single gene by region reference matrix of expression values.

### Regions of interest

In addition to brain-wide analyses that consider all regions, we focused on brain regions and circuits previously associated with anorexia nervosa and feeding behaviour. Specifically, we examined expression profiles of sites of deep brain stimulation studies: the nucleus accumbens and the subgenual region of the anterior cingulate cortex (Brodmann area 25) (Lipsman et al. 2017; Wu et al. 2013). The hypothalamus and ventral tegmental area provide two additional regions that have been named as potential sites (Nestler 2013). Guided by experimental mouse studies of feeding, we focused on several additional regions: arcuate nucleus of the hypothalamus, raphe magnus, raphe obscurus, amygdala, nucleus of the solitary tract and the parabrachial nucleus (Morton, Meek, and Schwartz 2014). To align the Allen fetal atlas with the regions of interest we merged six regions to form an expression profile for the subgenual cingulate cortex and two regions for the central amygdala.

### Brain region enrichment analysis

To calculate enrichment for the gene sets of interest within a brain region, z-scores in the processed expression matrices are ranked within a region, with high ranks marking specific expression. Our genes of interest were then projected into these ranked lists for each region. The area under the receiver operating curve (AUROC) statistic was used to quantify if the AN associated genes are more specifically expressed (ranked higher) in this sorted list of genes for a specific region or are randomly distributed within the sorted list. To assess statistical significance, the Wilcoxon Mann-Whitney U test was used. Benjamini-Hochberg false discovery rate procedure was used to correct for testing of multiple brain region tests within a dataset.

### Anorexia Nervosa associated gene lists

#### iPSC gene list

The initial discovery gene set contained 361 differentially expressed genes obtained from iPSC-derived neurons from AN patients compared to controls (Negraes et al, 2016).

Specifically, we used the gene symbols in Supplementary Table S5. To improve integration with the Allen data, we mapped antisense and intronic transcripts to the symbol of the protein coding gene by removing -AS and -IT suffixes. After these edits, 289 of 361 genes matched to the gene symbols of processed adult and fetal human brain gene expression datasets.

#### Whole exome sequencing lists

Genes containing rare variants associated with disordered eating as determined by exome sequencing were obtained from Lutter et al. (2017). Specifically, Table S3 provided a list of 186 genes that harboured damaging variants in individuals with the restricting type of anorexia nervosa and Table S4 provided a list of 245 damaging variants in individuals with binge-eating episodes which was used as a comparison gene set to test specificity. For both of these gene lists, we applied conservative significance threshold by calculating the Bonferroni corrected p-values with an estimate of 20,000 tested genes. After applying this filter, 51 genes associated with restricted eating and 80 genes associated to binge-purge eating disorders remained.

### Cell type-specific marker genes

Marker genes for major cell types were obtained from the NeuroExpresso database that performed a cross-laboratory analysis of several gene expression studies (Ogan et al. 2017). The marker sets were obtained from https://github.com/oganm/neuroExpressoAnalysis. In addition to the combined set, marker genes from cortical, brainstem, and amygdala analyses were used (based on the regions of interest).

### Gene expression responses to food deprivation in mice

Two mouse datasets were used to determine which of the AN associated genes are differentially expressed in response to food deprivation (Chen et al. 2017; Campbell et al. 2017). Both performed single-cell RNA sequencing to characterize transcriptional changes after food deprivation in the mouse hypothalamus.

The Chen et al. dataset assayed the transcriptome with single-cell RNA sequencing in over 14,000 cells from the mouse hypothalamus (adult female B6D2F1 mice) (Chen et al. 2017). The “Cells.Expresssion.Matrix.log_tpm+1” file from GSE87544 was used. This file contains expression values for 23,284 genes and 14,437 cells. We filtered for genes with human homologs in the homologene database, resulting in 16,155 genes (O’Leary et al. 2016). Genes with no expression in all cells were removed, leaving 15,217. Only cells from batch 1 and 2 were used because they contained data from both food deprived and control animals, leaving 10,983 cells. The food deprived animals were given only water for a 24 hour period.

The Campbell et al. dataset contains gene expression profiles of over 20,000 cells from the arcuate hypothalamus and median eminence (transgenic with C57BL6/J background)
(Campbell et al. 2017). Summarized gene expression profiles were obtained from the file named “GSE93374_Merged_all_020816_BatchCorrected_LNtransformed_doubletsremoved_Data.txt.g z”. This file contains expression values for 19,743 genes and 20,922 cells. To be consistent with the Chen dataset, we filtered these cells for those from female mice and only used batch 6 which also compared fasted (24 hours) to normal mice (ad libitum chow fed). After filtering out genes without human homologs and those without any variance in expression, 14,507 remained for 1,016 cells.

The Mann–Whitney U test was used to test for differential expression between the food deprived or hunger conditions. This method has been shown to be adequate for testing differential expression in single-cell studies with simple designs (Soneson and Robinson 2018). For each gene, the three dataset/batches were tested separately (13,208 mouse genes in both datasets). Fisher’s method was used to combine these three p-values into single meta p-values for up- and down-regulation of expression after 24 hours of food deprivation.

### Availability

Scripts, supplementary tables, and data files for reproducing the analyses are available online at https://github.com/derekhoward/molecularAN and https://figshare.com/s/72bfff921a5576f1a9e1.

## Results

Across the three AN associated gene sets, four genes overlap between the Lutter damaging variants and Negraes iPSC derived gene set (*TNFRSF10A*, *SCGB1A1*, *IFIT3*, and *GIPC3*; hypergeometric test, p < 0.02). In contrast, only one gene overlaps with the binge eating associated genes identified by Lutter et al. (*BDNF*). The six genes overlapping with the rs4622308 locus did not appear in the Lutter or Negraes sets. To characterize brain-wide neuroanatomical expression patterns we used the Negraes list because it is the largest, providing the most statistical power to detect regions with specific enrichment.

### Regional expression enrichment of the Negraes iPSC gene list

We first characterized the expression pattern of the genes that were differentially expressed in iPSC-derived neurons from AN patients in comparison to controls (Negraes et al. 2017). We used the adult and prenatal human gene expression data from the Allen Brain atlases to evaluate brain-wide expression patterns. For the 289 genes that remained after integration, our analysis identified 40 of 232 brain regions in the adult brain and 128 of 516 fetal brain regions where expression of the Negraes genes was higher than expected (all AUROC > 0.538, p_FDR_ < 0.05, Supplement tables 1 and 2). The top region in the adult analyses was the lateral parabrachial nucleus (AUROC = 0.579, p_FDR_ < 10^−4^), followed by the pontine raphe nucleus (AUROC = 0.574, p_FDR_ < 0.0005). Similarly, the lateral parabrachial nucleus and the raphe magnus nucleus rank second and third respectively in the fetal analyses (AUROC > 0.62, p_FDR_ < 10^−9^). Because these regions have been previously associated with feeding behaviour in mice and rats, we focused our analyses on several regions of interest. These regions were selected from circuits previously associated with feeding in rodents or have been targeted by deep brain stimulation for the treatment of AN (or proposed targets). In the adult expression data, 4 of the 10 regions of interest preferentially expressed the Negraes genes after correction for the 232 tested regions (pontine raphe nucleus, ventral tegmental area, lateral and medial parabrachial nucleus; all p_FDR_ < 0.0005; Figure 2). In the fetal brain, these four regions and an additional three regions of interest had enriched expression after correction for the 516 tested regions (all p_FDR_ < 0.02, Figure 3). One region, the subgenual cingulate cortex, was depleted for expression (AUROC = 0.425, p_FDR_ < 0.0001).

**Figure 2.**
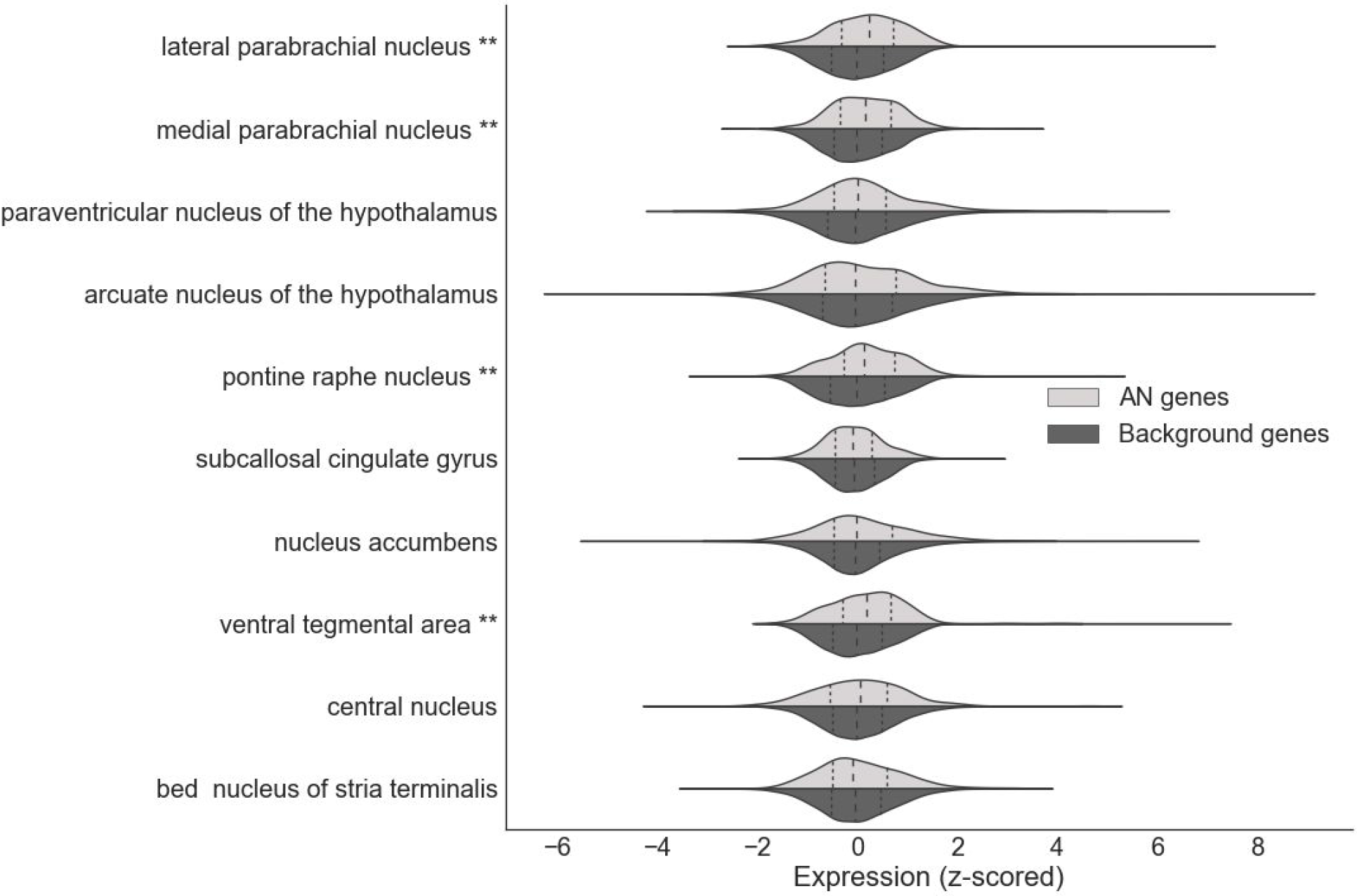
Violin plots showing gene expression patterns of the set of AN associated genes derived from Negraes et al. (light grey) with all remaining background genes shown in dark grey for the regions of interest in adult brain data. Vertical lines mark expression quartiles; ** denotes p_FDR_ < 0.05 from Mann-Whitney U tests.

**Figure 3.**
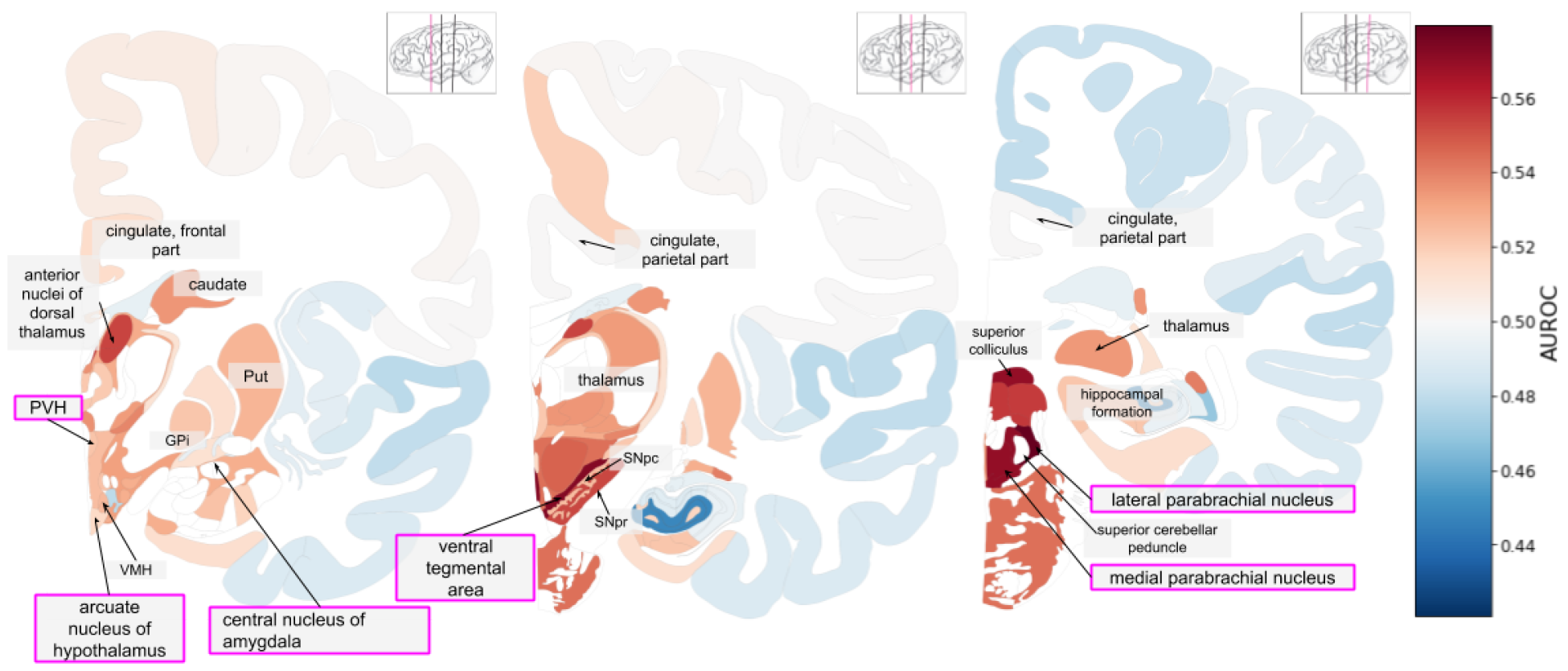
Anatomical maps showing aggregate gene expression patterns of the AN associated genes Negraes et al. in the adult brain. Regions of interest are highlighted with purple boxes. The inset thumbnail marks the current slice in pink. AUROC values range from depleted expression in dark blue to enriched in dark red with missing values in white. (PVH: paraventricular nucleus of hypothalamus; GPi: globus pallidus, internal segment; Put: putamen; VMH: ventromedial hypothalamic nucleus; SNpr: substantia nigra pars reticulata; SNpc: substantia nigra pars compacta)

**Figure 4.**
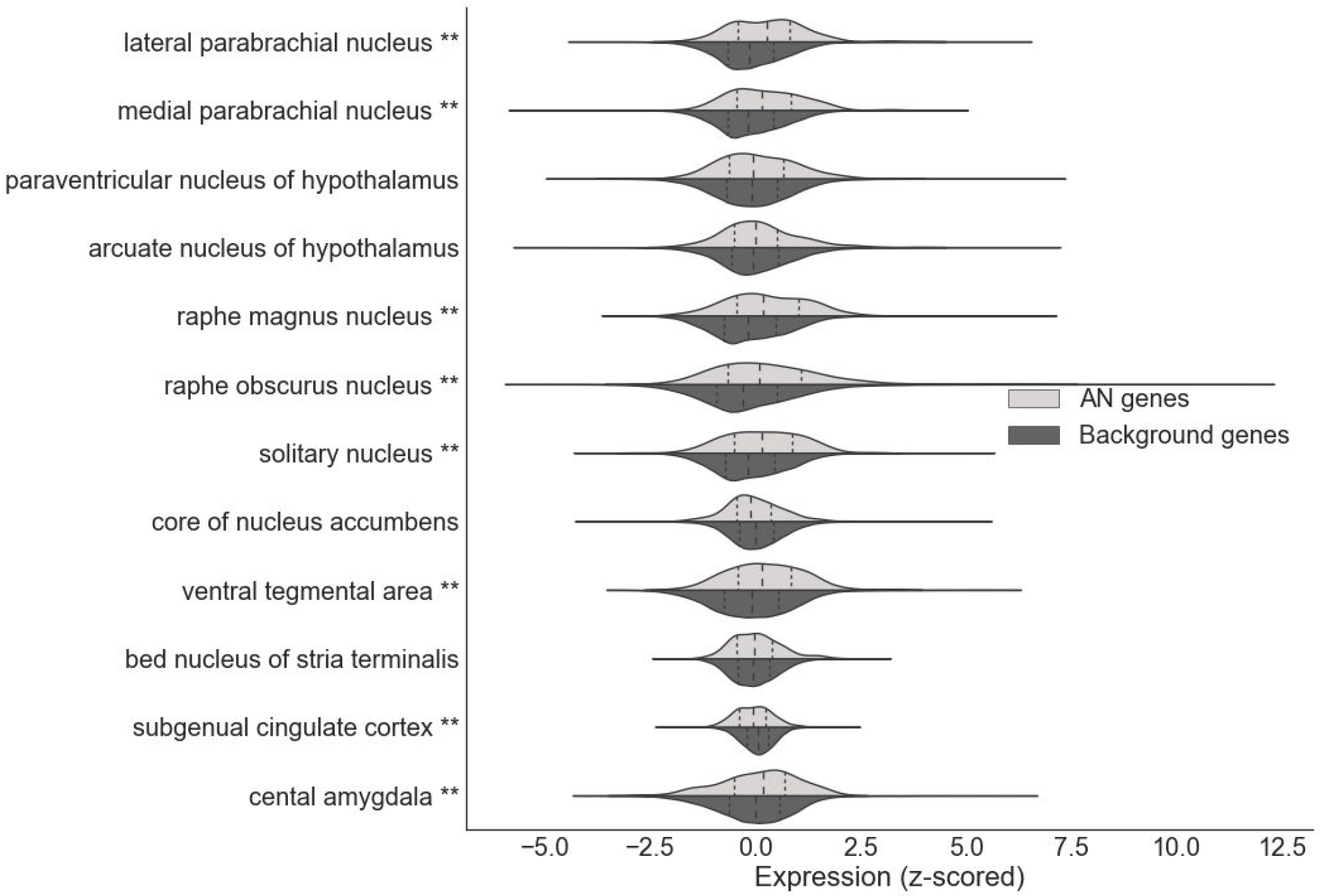
Violin plots showing gene expression patterns of the AN associated genes from Negraes et al. (light grey) with all remaining background genes shown in dark grey for the regions of interest in fetal brain data. Vertical lines mark expression quartiles; ** denotes p_FDR_ < 0.05 from Mann-Whitney U tests.

**Figure 5.**
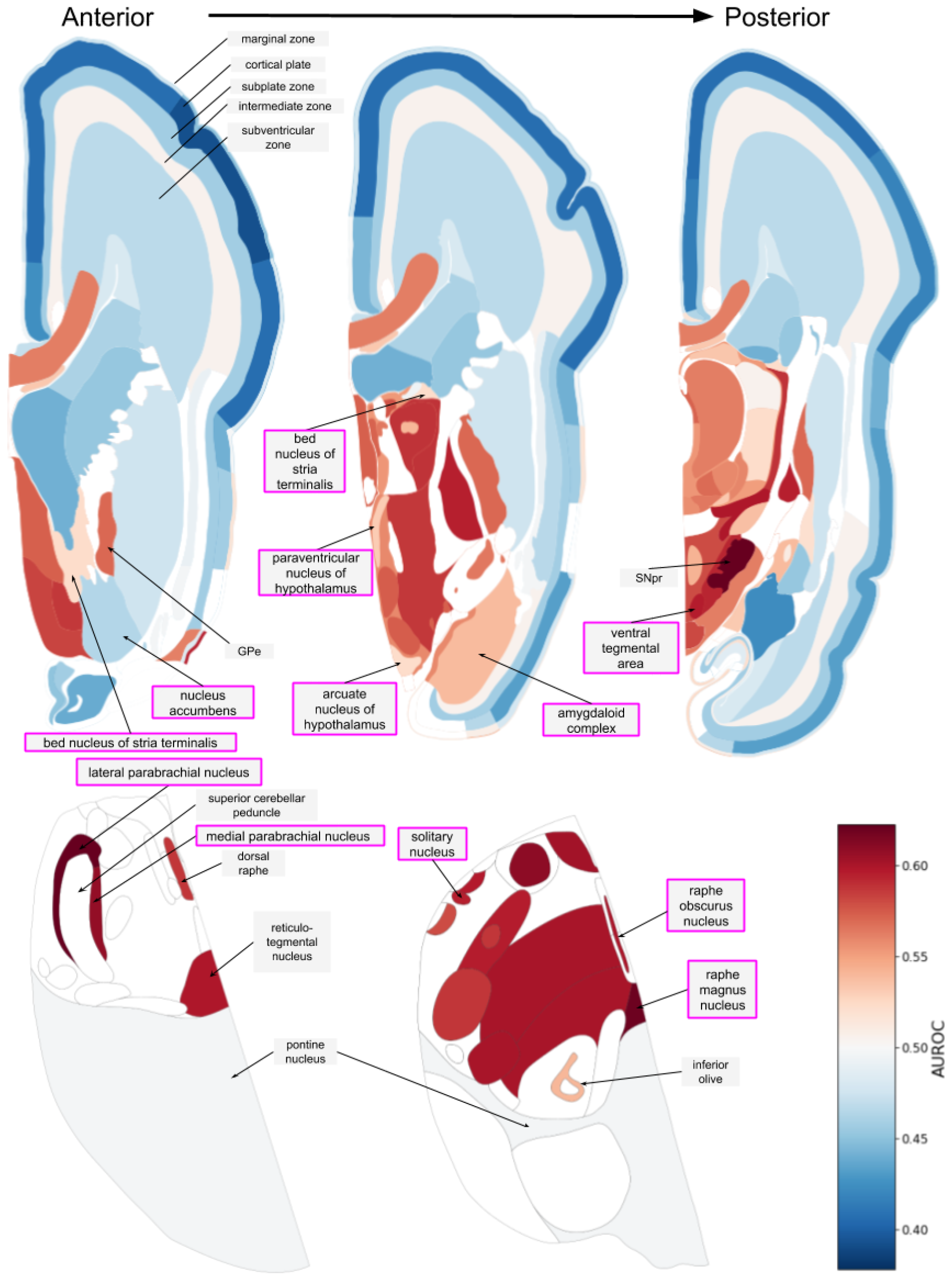
Anatomical maps showing aggregate gene expression patterns of the AN associated genes Negraes et al. in the fetal brain. Regions of interest are highlighted with purple boxes. AUROC values range from depleted expression in dark blue to enriched in dark red with missing values in white. (GPe: globus pallidus external segment; SNpr: substantia nigra pars reticulata)

### Regional expression enrichment of the Duncan GWAS gene list

To provide an independent test of the regions identified from the Negraes list, we examined genes from the largest genome-wide association study (GWAS) of anorexia nervosa to date that observed a single locus (rs4622308) with genome-wide significance. This locus overlaps with six nearby genes (*IKZF4*, *RPS26*, *ERBB3*, *PA2G4*, *RPL41*, and *ZC3H10*) (Duncan et al. 2017). Unlike the above brain-wide analysis, we only tested for expression enrichment of these genes in the regions of interest. In the adult data, specific expression was observed in the lateral parabrachial nucleus (AUROC = 0.74, uncorrected p < 0.03) and the ventral tegmental area (AUROC = 0.69, uncorrected p < 0.06). In the fetal data, specific expression was observed in the arcuate nucleus of the hypothalamus (AUROC = 0.71, uncorrected p < 0.05) and the ventral tegmental area (AUROC = 0.7, uncorrected p < 0.05). Expression rankings of the six genes in these regions are marked in Figure 4. Of these three regions, the ventral tegmental area and lateral parabrachial nucleus were also brain-wide significant when using the Negraes list, increasing the confidence of their involvement in AN.

**Figure 6.**
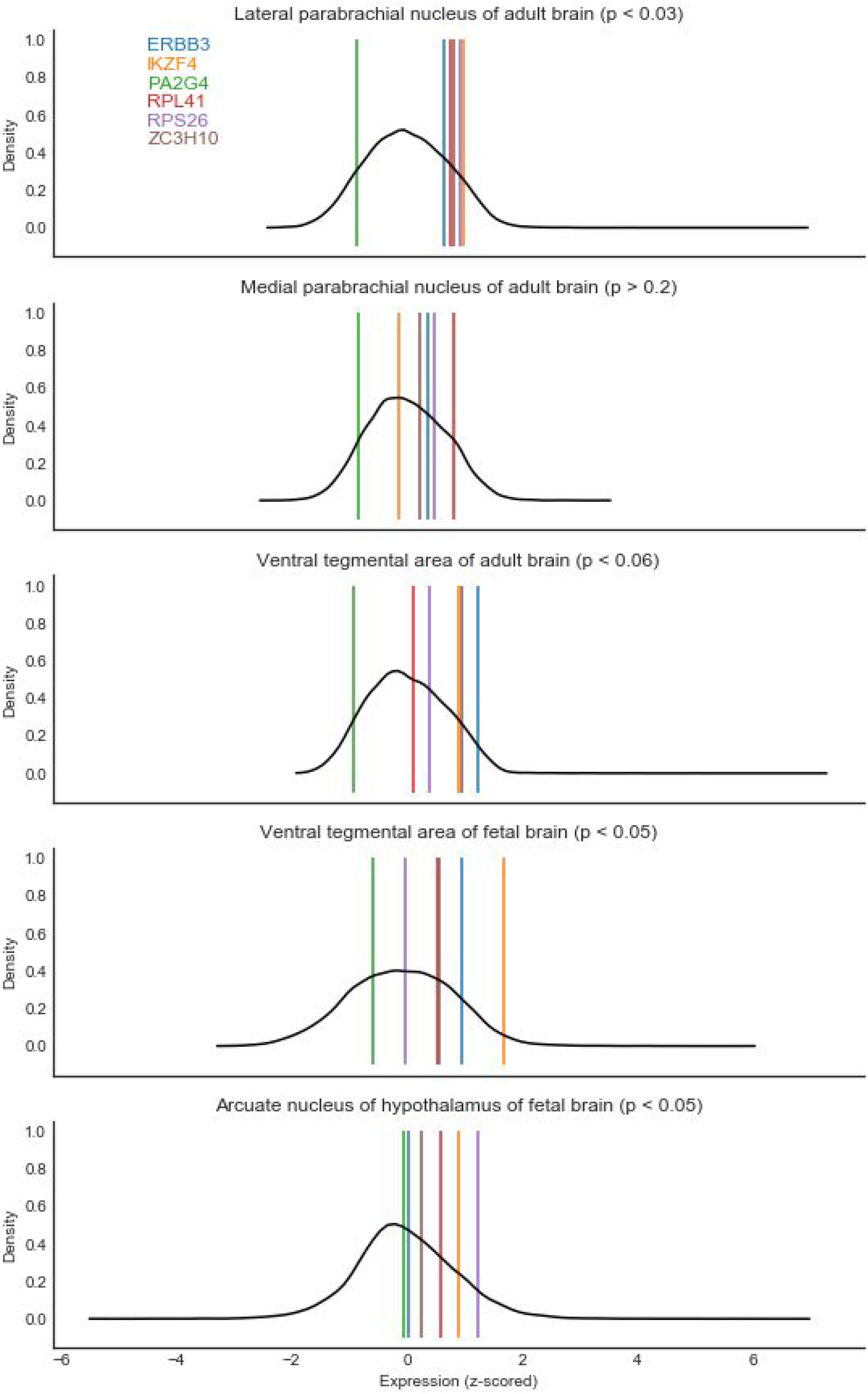
Density plots of z-scored genome-wide expression within a brain region for either the adult or fetal brain reference atlases (in black). Coloured lines mark expression of the 6 genes near rs4622308.

### Regional expression enrichment of the Lutter gene variants

To further test our findings of regionally enriched expression we assessed the patterns of genes that harbour novel and ultra-rare damaging variants in anorexia nervosa cases (restricting subtype) (Lutter et al. 2017). As shown in Table 1, specific expression was observed in the bed nucleus of the stria terminalis (AUROC = 0.6, uncorrected p < 0.02) and the lateral parabrachial nucleus in the adult brain data (AUROC = 0.58, uncorrected p < 0.05). In the fetal brain, no regions reached significance but all except the subgenual cingulate cortex had higher expression than average (AUROC > 0.5). We also note that the lateral parabrachial nucleus had the highest AUROC value of the 13 fetal regions of interest tested (AUROC = 0.566, uncorrected p < 0.11). Using the second list of genes provided by Lutter et al. that were found to harbour damaging variants in in cases with binge-eating behaviour were not significantly enriched in any of the brain regions of interest in the adult or fetal brains (obtained from patients with AN-binge/purge subtype, bulimia nervosa, and binge eating disorder diagnoses). In contrast, they were significantly depleted in the medial parabrachial nucleus, nucleus accumbens, and the arcuate nucleus of the hypothalamus (all AUROC < 0.44, uncorrected p < 0.05). Overall, this further highlights the lateral parabrachial and suggests subtype differences.

**Table 1.** Region of interest focused expression enrichment results for genes that harbour damaging variants in anorexia nervosa cases (restricting subtype) using adult expression data.

| Brain region | AUC | p |
| --- | --- | --- |
| lateral parabrachial nucleus | 0.580 | 0.0477 |
| medial parabrachial nucleus | 0.577 | 0.0572 |
| paraventricular nucleus of the hypothalamus | 0.524 | 0.549 |
| arcuate nucleus of the hypothalamus | 0.487 | 0.751 |
| pontine raphe nucleus | 0.554 | 0.181 |
| subcallosal cingulate gyrus | 0.471 | 0.470 |
| nucleus accumbens | 0.579 | 0.0508 |
| ventral tegmental area | 0.570 | 0.0855 |
| central nucleus | 0.555 | 0.176 |
| bed nucleus of stria terminalis | 0.601 | 0.0123 |

### Regional expression enrichment summary

Our results across the three sources of AN genes and two anatomical expression atlases are summarized in Tables 2 and 3. When combined, enrichment in the lateral parabrachial nucleus is consistently found in the adult atlas. In the fetal brain, higher expression in the ventral tegmental area is observed for two of the anorexia associated gene sets. Regional expression profiles for each of targetted are available in Supplemental table 3.

**Table 2.** Summary of expression enrichment in the adult brain expression atlas (** denotes p_FDR_ < 0.05, + denotes p < 0.05, and - marks non-significant enrichment).

| Brain region | Negraes | Duncan | Lutter restricted-eating | Lutter binge-eating |
| --- | --- | --- | --- | --- |
| lateral parabrachial nucleus | ** | + | + | - |
| medial parabrachial nucleus | ** | - | - | - |
| paraventricular nucleus of the hypothalamus | - | - | - | - |
| arcuate nucleus of the hypothalamus | - | - | - | - |
| pontine raphe nucleus | ** | - | - | - |
| subcallosal cingulate gyrus | - | - | - | - |
| nucleus accumbens | - | - | - | - |
| ventral tegmental area | ** | - | - | - |
| central nucleus | - | - | - | - |
| bed nucleus of stria terminalis | - | - | + | - |

**Table 3.**
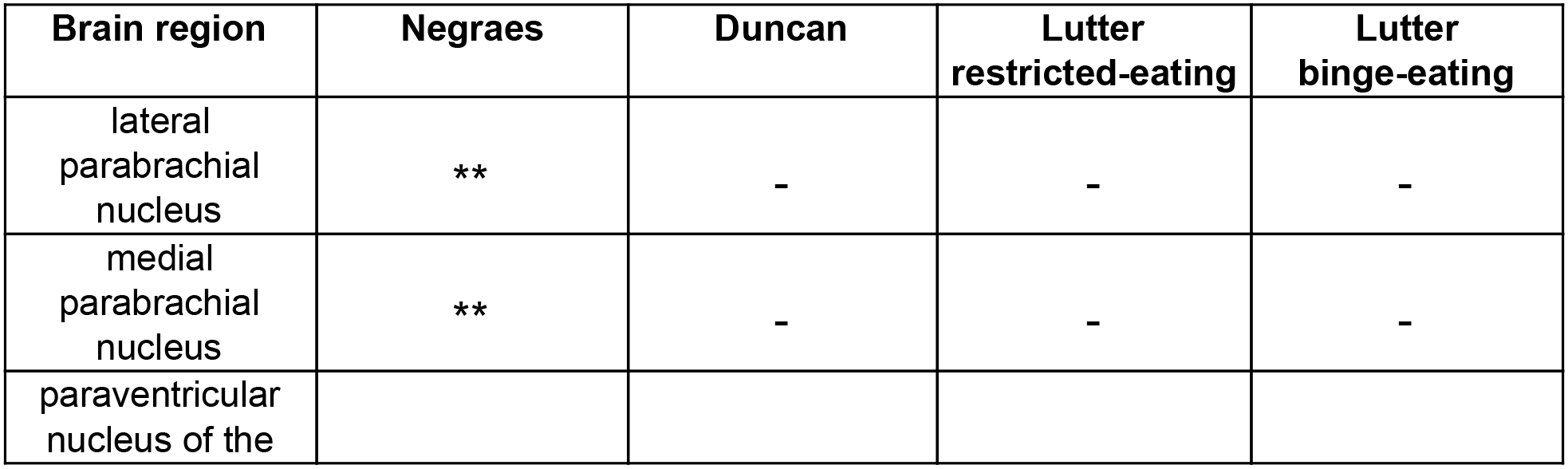
Summary of expression enrichment in the fetal brain expression atlas (** denotes p_FDR_ < 0.05, + denotes p < 0.05, and - marks non-significant enrichment).

**Table 3.**
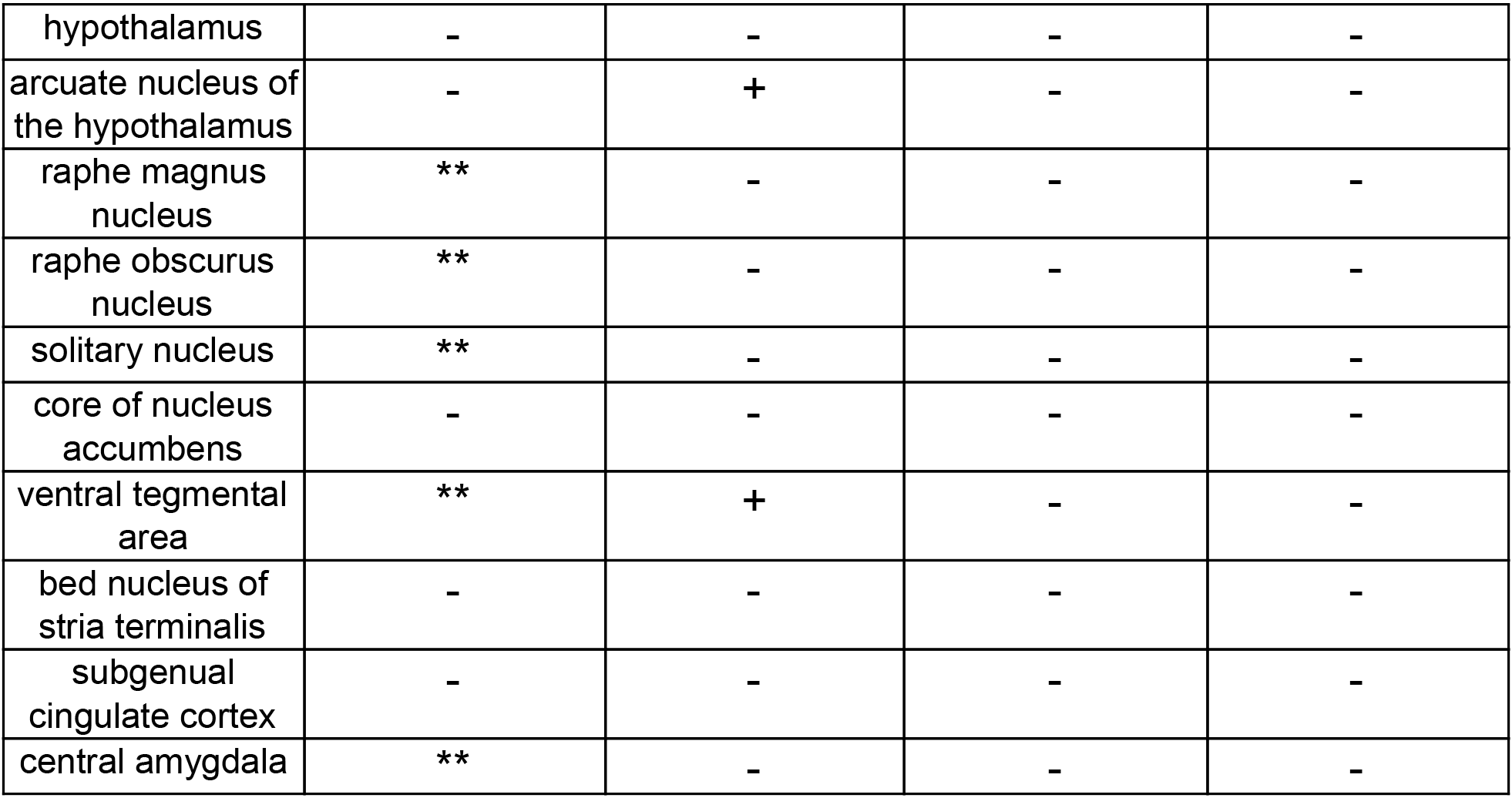

### Cell type marker enrichment

We used the NeuroExpresso database to determine if the gene lists are enriched for cell type markers. Only the Negraes gene set had significant cell type marker enrichment. Specifically, between 8 and 9 genes overlapped with the microglia markers across the four regional lists (all, cortex, amygdala, and brainstem). While the Neuroexpresso marker genes were obtained from analyses of whole brain homogenates, the microglia marker genes from the brainstem lists had the lowest corrected p-value (9 of 137 genes overlap, hypergeometric test, p_FDR_ < 0.01). These genes are split between up- and down-regulated in the Negraes results, suggesting the signal is not due to different microglia proportions (5 down-regulated and 4 up-regulated, Supplement Table 4). While not significant, overlap with the separate lists of microglia activation and deactivation were 4 and 2 genes respectively. In addition, the largest overlap for the Lutter gene list was with the brainstem microglia activation markers (*IL17RA* and *SLA;* uncorrected p < 0.05). In the Duncan set, only the *ERBB3* gene was a Neuroexpresso marker (for oligodendrocyte precursors in the cortex analyses). The binge eating associated genes from the Lutter study did not show any clear enrichment (1-2 genes per cell type), however, we note that *PHOX2A* was contained in the small list of noradrenergic markers. Overall, we find more than the expected number of microglia marker genes in two of the three AN associated gene sets.

### Gene expression responses to food deprivation in mice

We next used two single-cell studies to determine if the genes of interest are differentially expressed as a result of fasting in the female mouse hypothalamus (11,999 total cells) (Campbell et al. 2017; Chen et al. 2017). Genome-wide, 2,160 of the 13,208 genes we examined were significantly differentially expressed (16.3%, p_FDR_ < 0.05). None of the gene lists of interest were disproportionately enriched for these genes (17.6 to 20.6%). While not significantly enriched, genes that are altered after fasting and associated with anorexia nervosa are of interest (full list provided in Supplement Table 5). For example, in the Duncan set, Rps26 was significantly down-regulated after fasting in both datasets (Figure z, meta-p_FDR_ < 10^−13^). However, this finding is not consistent with one batch of the Chen dataset showing up-regulation after fasting (meta-p_FDR_ < 10^−4^). In contrast, we did not detect any significant changes in expression for the other four genes with mouse homologs near rs4622308.

**Figure 7.**
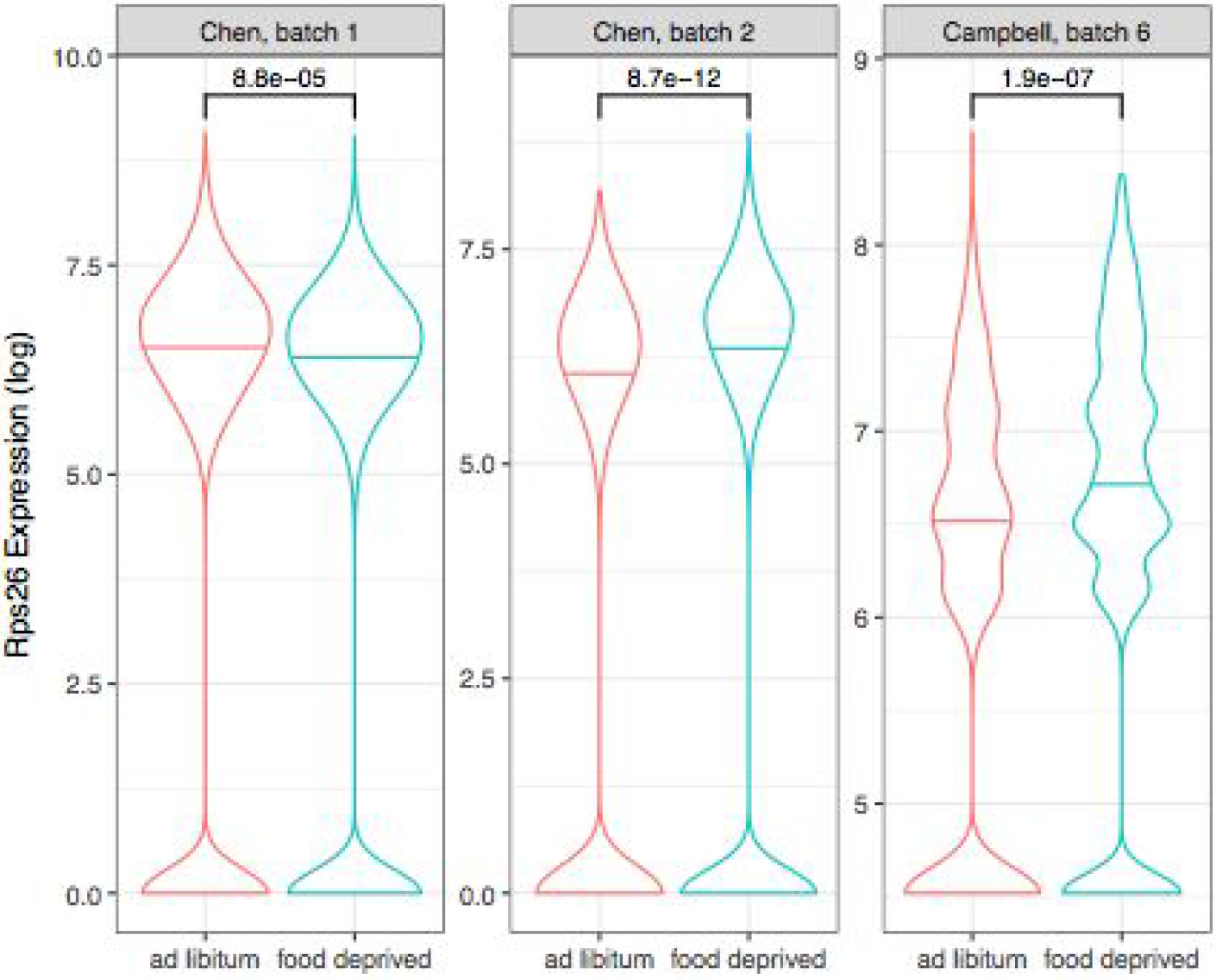
Violin plots of Rps26 expression in *ad libitum* (red) and food deprived (blue) conditions. The first two plots are from the Chen dataset where expression is measured by the log of transcripts per million. The last plot is from the Campbell dataset, which measured expression by the natural log of the counts per million plus one (unique molecular identifier method). Connecting brackets at the top provide p-values from Mann-Whitney U tests.

## Discussion

We first investigated the neuroanatomical expression patterns of genes associated with anorexia nervosa. Our results show that AN associated genes are highly expressed in regions linked to food intake and reward in rodent studies, suggesting direct relevance to human studies of anorexia nervosa. The most consistent region of enrichment is the parabrachial nucleus. In rodents, this region was once named the “pontine taste area” (Saper 2016) and plays a key role in appetite regulation. For example, calcitonin gene-related peptide (CGRP) neurons in the lateral parabrachial nucleus have been shown to inhibit feeding, suppress appetite (Carter et al. 2013), participate in conditioned taste aversion (Reilly 1999), control meal termination (Campos et al. 2016), and prevent overeating (Campos et al. 2018). More broadly, CGRP neurons in the parabrachial are thought to serve as a general purpose alarm that encodes diverse danger signals (Saper 2016; Palmiter 2018). The *CALCB* gene, which encodes the beta isoform of CGRP is the 23rd most differentially expressed gene from the AN derived stem cell study (Negraes et al. 2017). Within the AN associated genes we studied, *CALCB* has the most specific expression in the medial parabrachial nucleus in both the adult and fetal human brain data and ranks 2nd and 4th in the fetal and adult lateral parabrachial nuclei respectively (Supplement Table 3). Within the Negraes list, we also note that *RAMP1*, a CGRP receptor transporter is specifically expressed in regions receiving parabrachial inputs such as the bed nucleus of the stria terminalis (ranked 11/342), the paraventricular nucleus of the hypothalamus (ranked 13/342) and the central amygdala (ranked 16/342). Mouse studies have also revealed that the lateral parabrachial contains thermosensory relay neurons and is involved in thermoregulation (Geerling et al. 2016; Yahiro et al. 2017), suggesting relevance of this region to the colder body surface temperatures observed in anorexia nervosa patients (Belizer and Vagedes 2018; Wakeling and Russell 1970). Our results also point to the ventral tegmental area which receives hypothalamic input from orexin neurons that have been linked to appetite and reward (Aston-Jones et al. 2010). Activation of orexin receptors in the ventral tegmental area promoted food intake in a hedonic feeding model (Terrill et al. 2016). While our results do not suggest disturbances in the hypothalamus or the bed nucleus of stria terminalis, they do link anorexia nervosa to subcortical appetitive circuits.

Past trials of deep brain stimulation for severe and treatment-resistant anorexia nervosa have targeted the subcallosal cingulate and the nucleus accumbens (Lipsman et al. 2017; Nestler 2013). These sites were chosen based on results from MRI studies that were limited to the cortex and ‘top-down’ networks related to emotional homeostasis (Lipsman, Woodside, and Lozano 2015). We did not observe expression enrichment of AN associated genes in the subcallosal cingulate or nucleus accumbens. However, our results support targeting of the ventral tegmental area which has been recently been proposed as a target site (Nestler 2013). In addition, our findings suggest that stimulation of the parabrachial nucleus in the pontine tegmentum may be effective in severe treatment resistant cases. While there is a high risk of complications from brainstem surgery, we note that the pontine tegmentum has been a safe target of deep brain stimulation for over a decade (Mazzone et al. 2016).

While we note that two CGRP related genes are highly ranked, at a broader cell-type level we found enrichment of microglia genes. This is found first in the Negraes stem cell derived genes and, to a lesser degree, in the list of damaging variants that were associated with AN. In mice, stimulation of the innate immune system by activation of toll-like receptor 2 resulted in sickness behavior that included anorexia (Jin et al. 2016) and aberrant agouti-related protein signalling in an anorexia mouse model was associated with microglial activation (Nilsson et al. 2008). Linking the regional results, the lateral parabrachial and the ventral tegmental area are enriched for the Neuroexpresso microglia marker genes (adult expression data: AUROC > 0.62, p_FDR_ < 10^−5^). Relative to the other regions, the lateral parabrachial nor the ventral tegmental area rank in the top 20 regions that are most enriched for microglia markers in adult expression data. In contrast, these two regions rank in the top 5 when using the Negraes stem cell derived genes, showing that the microglia enrichment cannot fully explain the regional results. Microglia are key contributors to sex differences in the brain from both structural and functional perspectives (Lenz and McCarthy 2015). In addition, microglial sex differences have been linked to behavioural differences and pain hypersensitivity (Sorge et al. 2015; Salter and Stevens 2017; Lenz et al. 2013). In summary, our results suggest microglia deserve more attention in AN research and that they may help explain the higher prevalence of AN in females.

Our use of transcriptomic and genetic resources that crosses species and development marked robust signals but limits finer interpretations. Developmentally, we observed regional enrichment in the prenatal and adult brain but lack expression data from the adolescent brain for our regions of interest. For diagnostic criteria, we found a subtype dependent signal when using the genes harbouring damaging variants that Lutter et al. identified (restricted eating vs. binge-eating). The other two sources of AN associated genes were not specific to AN subtypes, limiting our ability to detail this difference. Our main gene list was from the Negraes et al. stem cell study. Their differentiation procedure generated primarily cortical neurons, with a low proportion of glial cells. Our findings of microglia and subcortical expression enrichment suggest follow-up study of the Negraes et al. differentially expressed genes should include microglia and subcortical regions. While providing a weaker signal, we validated enrichment of the microglia markers and the lateral parabrachial and ventral tegmental areas with genetically associated genes that are not inherently linked to a specific tissue or cell type.

Within the stem cell derived gene list from Negraes et al., *TACR1* was identified as a novel an potential contributor to AN pathophysiology (2017). While lower in the fetal brain, in the adult data *TACR1* has high expression in the lateral parabrachial nucleus (ranked 22^nd^ of 232 regions). In relation to the other AN associated genes, *TACR1* expression ranks 56^th^ of 342 in the adult lateral parabrachial nucleus. When the three tachykinin related genes in the Negraes et al. are combined (*TAC1*, *TACR1*, and *TACR2*), the lateral parabrachial nucleus is enriched in the adult data (ranked 7th of 232 regions, AUROC = 0.88, uncorrected p < 0.02). This suggests that the proposed participation of the tachykinin system in AN involves the lateral parabrachial area.

Comparable transcriptomic studies of AN are limited. A post-mortem study by Jaffe et al. identified six differentially expressed genes in the prefrontal cortex of cases diagnosed with eating disorders (AN or bulimia nervosa) in comparison to controls (2014). While these six genes are not enriched in the lateral parabrachial or ventral tegmental areas, the most significant gene, *RFNG*, is strongly expressed in the adult lateral parabrachial nucleus (brain-wide: ranked 1st of 232 regions; genome-wide: 218 of 20,869 genes). The lack of more agreement with this study may be due to the mix of diagnoses, focus on the prefrontal cortex, or the effects of chronic illness in these post-mortem samples. Predicting gene expression from genetic information avoids these pitfalls by using reference expression data from healthy subjects. This transcriptomic imputation approach was recently applied to the genetic data used in the Duncan et al. GWAS (Huckins et al. 2018). Within the rs4622308 locus, they identified 35 associations where differential expression was predicted for a specific gene and tissue. Within the six genes we used to represent rs4622308, they found significant associations for only *RPS26*. They also found that predicted expression of *RPS26* was negatively correlated with BMI, weight, and waist circumference. In agreement, within these six genes, we also highlighted *Rps26* due to its significant down-regulated in the hypothalamus of food deprived mice. Unlike our study, Huckins et al. didn’t note enrichment in a specific brain region. This is probably due to the coarse resolution of the transcriptomic data that was limited to 13 large brain regions which did not include the lateral parabrachial or ventral tegmental areas. Within our regions of interest, they did test predicted expression for the hypothalamus and the anterior cingulate cortex but like us, they did not implicate these regions. While our results cannot be directly compared to past transcriptomic studies of cases and controls, they can be linked to the lateral parabrachial nucleus and also mark *RPS26* as a key gene in AN.

## Conclusion

In summary, we found that genes associated with AN are expressed at higher levels in the lateral parabrachial nucleus and the ventral tegmental area in comparison to the rest of the healthy adult and fetal brain. The adult expression enrichment of the lateral parabrachial is confirmed with genes from two independent genetic studies (Table 2). In the fetal brain, enrichment of the ventral tegmental area is also observed for the six genes near the only common variant associated with the disorder (Table 3). We also observed signals in the adult and fetal pontine raphe, but they were not repeated when using the genes from the genetic studies. We also found more than the expected number of microglia marker genes in the AN associated genes. Finally, using mouse transcriptomic data, we noted several of the AN associated genes are differentially expressed during food deprivation. While these genes that respond to fasting are not enriched in the AN associated gene sets, we highlight Rps26 which is proximal to the only common variant associated with AN.

## Acknowledgements

We thank the Allen Institute for Brain Science and the BrainSpan Consortium Members for creating the reference transcriptomic atlases of the fetal and adult brain.

This study was supported by the CAMH Foundation and a National Science and Engineering Research Council of Canada (NSERC) Discovery Grant to LF.

